# Four-color single-molecule imaging system for tracking GPCR dynamics with fluorescent HiBiT peptide

**DOI:** 10.1101/2024.06.10.598203

**Authors:** Toshiki Yoda, Yasushi Sako, Asuka Inoue, Masataka Yanagawa

## Abstract

Single-molecule imaging provides information on diffusion dynamics, oligomerization, and protein-protein interactions in living cells. To simultaneously monitor different types of proteins at the single-molecule level, orthogonal fluorescent labeling methods with different photostable dyes are required. G-protein-coupled receptors (GPCRs), a major class of drug targets, are prototypical membrane receptors that have been studied using single-molecule imaging techniques. Here we developed a method for labeling cell-surface GPCRs inspired by the HiBiT system, which utilizes the high affinity complementation between LgBiT and HiBiT fragments of the NanoLuc luciferase. We synthesized four fluorescence-labeled HiBiT peptides (F-FiBiTs) with a different color dye (Setau-488, TMR, SaraFluor 650 and SaraFluor 720). We constructed a multicolor total internal reflection fluorescence microscopy system that allows us to track four color dyes simultaneously. As a proof-of-concept experiment, we labeled an N-terminally LgBiT-fused GPCR (Lg-GPCR) with a mixture of the four F-FiBiTs and successfully tracked each dye within a cell at the single molecule level. The F-FiBiT-labeled Lg-GPCRs showed agonist-dependent changes in the diffusion dynamics and accumulation into the clathrin-coated pits as observed with a conventional method using a C-terminally HaloTag-fused GPCR. Taking advantage of luciferase complementation by the F-FiBiT and Lg-GPCRs, the F-FiBiT was also applicable to bioluminescence plate-reader-based assays. By combining existing labeling methods such as HaloTag, SNAP-tag, and fluorescent proteins, the F-FiBiT method will be useful for multicolor single-molecule imaging and will enhance our understanding of GPCR signaling at the single-molecule level.

## Introduction

Total internal reflection fluorescence microscopy (TIRFM) enables real-time observation of single molecules with sparse fluorescent labeling techniques. Single-molecule tracking (SMT) analysis provides quantitative insights into parameters such as oligomer size distribution, diffusion dynamics, and spatial distribution of fluorescently labeled molecules at the plasma membrane. Extension of single-molecule imaging analysis to multi-color allows estimation of association and dissociation rates of protein-protein interactions in living cells [1]. Single-molecule microscopy has been instrumental in assessing the dynamics of membrane receptors, transducers, and lipid probes, leveraging its unique capability to quantify multiple dynamics parameters in a single measurement, which are otherwise challenging to measure in living cells. This approach aids in unraveling the intricate spatiotemporal regulatory mechanisms of membrane receptor signaling that govern various cellular responses [2,3].

Earlier studies using fluorescent ligands have shown that membrane receptors such as receptor tyrosine kinases (RTKs) and G-protein-coupled receptors (GPCRs) are in a dynamic monomer-dimer equilibrium [4–7]. The dimerization and oligomerization are essential for the phosphorylation and signaling of RTKs, including ErbB family receptors. The kinetic analysis of ligand binding to ErbB receptors at the plasma membrane allowed to distinguish between on- and off-rates to the receptors in a different oligomeric state, predicting the presence of the predimer and high affinity intermediates prior to the agonist binding to each protomer [4,5]. The monomer-dimer transition rates of GPCRs, including the M1 muscarinic receptor [6] and the N-formyl peptide receptor [7], were quantified in a similar manner, showing that class A GPCRs are in a dynamic equilibrium state and do not form a stable dimer as assumed from bulk measurements. However, the fluorescent ligand approach does not provide information on receptor dynamics in the apo-state before ligand stimulation.

Fluorescent protein fusion has become a preferred method for tracking membrane protein dynamics, effectively resolving the restriction of the fluorescent ligand approach, but it still has some limitations. For instance, mEGFP [8], one of the gold standard fluorescent proteins used in single-molecule imaging, has lower brightness and photostability compared to organic fluorescent dyes developed for super-resolution microscopy, such as Janeria Fluor [9] and SaraFluor dyes [10]. This limitation hampers long-term single-molecule measurements over a few seconds.

Furthermore, since all expressed fusion proteins emit fluorescence, it is imperative to attenuate their expression levels to a density (∼1 **μ**m^-2^) conducive to single-molecule imaging, which necessitates careful adjustment of transfection conditions [1].

The development of fluorescent tag systems, such as SNAP-tag and HaloTag, has overcome the limitations of the fluorescent ligand and fusion protein approaches [11,12]. The HaloTag system offers a range of bright and photostable commercially available fluorescent dyes compared to SNAP-tag. Considering that the apparent affinity to bind the dye is one order of magnitude higher than that of the SNAP-tag, the HaloTag would be the preferred choice for single-molecule imaging. However, due to the larger size of HaloTag (∼33 kDa) compared to SNAP-tag (∼20 kDa), HaloTag fusion sometimes inhibits the function of target proteins. These tagging systems allow comparison of membrane receptor behavior before and after ligand stimulation.

Through the combination of fluorescent labeling techniques and the use of multiple color dyes, it becomes possible to perform two- or three-color simultaneous single-molecule imaging. The multicolor single-molecule imaging has been used to observe the interaction of membrane receptors and transducers. Studies that simultaneously observed GPCRs and G proteins discovered a membrane domain where the two molecules accumulate and actively transduce signals [13,14]. The dynamic interaction between GPCRs and GPCR kinase (GRK) or β-arrestins are also observed using two-color single-molecule imaging, suggesting that the co-accumulation of GPCR and transducers in a nanodomain plays an important role for signaling attenuation or biased signaling [15,16]. For many GPCRs, agonist stimulation causes an increase in the fraction in the immobile nanodomain, reflecting GRK-dependent phosphorylation [16], recruitment of β-arrestin [18,19], and subsequent accumulation in clathrin-coated pits (CCP) [17].

To achieve simultaneous single-molecule imaging of membrane receptors and various signaling molecules, an additional fluorescent labeling technique is required. In the present study we develop a method for labeling cell surface receptors inspired by a split luciferase system [20]. The HiBiT system utilizes the complementarity of two NanoLuc fragments called Large BiT (LgBiT, ∼18 kDa) and HiBiT peptide (∼1.3 kDa). Conventionally, the cell surface expression of HiBiT-fused GPCRs has been monitored by bioluminescence from reconstituted NanoLuc in the presence of high affinity HiBiT peptide and furimazine substrate [21,22]. We synthesized fluorescently-labeled Flag-HiBiT (F-FiBiT) peptide to label LgBiT-fused GPCRs with various color dyes to achieve four-color simultaneous single-molecule imaging. As shown previously using C-terminally HaloTag-fused GPCR, we observed agonist-dependent changes in the diffusion dynamics of GPCR and its accumulation in CCP by F-FiBiT approach for LgBiT-fused GPCR (Lg-GPCR) [17]. F-FiBiT was not only applicable for the single-molecule imaging but also for the flow cytometry and bioluminescence-based plate reader assay, which expanded our experimental options with a single plasmid construct.

## Materials and Methods

### Materials

TMR-labeled Flag-HiBiT (TMR-FiBiT: TMR-DYKDDDDKGDGSVSGWRLFKKIS) and TMR-labeled HiBiT (TMR-HiBiT: TMR-VSGWRLFKKIS) were synthesized in Genscript. SeTau-488-NHS (SETA BioMedicals)-labeled Flag-HiBiT (ST488-FiBiT), SaraFluor650-NHS (GORYO CHEMICAL)-labeled Flag-HiBiT (SF650-FiBiT) and SraFluor720-NHS (GORYO CHEMICAL)-labeled Flag-HiBiT (SF720-FiBiT) were synthesized and purified in RIKEN CBS, RRD peptide synthesis service. F-FiBiT were dissolved in dimethyl sulfoxide (DMSO, Wako) and used as stock solutions. The concentration of the stock solutions was quantified based on the absorbance of the F-FiBiT diluted in PBS.

### Plasmid constructions

For Lg-GPCR construction, GPCR N-terminally fused EGFR-signal-sequence, linker (5’-ATGCGACCCTCCGGGACGGCCGGGGCAGCGCTCCTGGCGCTGCTGGCTGCGCTCTGCCCGGCGAGTCGGGCTACCCGCTTAAAAGCTTGGCAATCCGGTACTGTTGGTAAAGCCACC-3’), LgBiT and linker (5’-GGGAGTTCCGGTGGTGGCGGGAGCGGAGGTGGAGGCTCGAGCGGT-3’) inserted into the HaloTag-removed-pFC15A vector (Promega). GPCR sequences were obtained from PRESTO-tango kit (Addgene Kit #1000000068). For the construction of SNAP-tag fused CD86, EGFR-signal-sequence, linker (5’-ACCCGCTTAAAAGCTTGGCAATCCGGTACTGTTGGTAAAGCCACC-3’), SNAP-tag, linker (5’-GGGAGTTCCGGTGGTGGCGGGAGCGGAGGTGGAGGCTCGAGCGGT-3’), and CD86 (1-277) were inserted into the HaloTag-removed-pFC15A vector in the above order from the N-terminal end. LgBiT-fused CD86 (Lg-CD86) and HaloTag-fused CD86 (Halo-CD86) were constructed by swapping SNAP-tag sequence on the SNAP-CD86 with LgBiT and HaloTag sequence, respectively. In the construction of the above plasmid DNA, all DNA fragments were assembled using a seamless cloning method [25,26]. The cDNA of EGFP-tagged clathrin-light-chain (EGFP-CLC) [25] and C-terminally EGFP-fused CD86 (CD86-EGFP) were constructed as previously reported [17]. The HiBiT-V2R/pCAGGS (Hi-V2R) was constructed by inserting the sequence, HiBiT-linker (TCAGGGGGGTCCGGGGGGGGCGGATCAGGCCTC)-V2R, into a multi cloning site (MCS) of pCAGGS by NEBuilder. The β-arrsstin2/pCAGGS was constructed by inserting the β-arrsstin2 (human) sequence into MCS of pCAGGS by NEBuilder.

### Cell culture

Cell culture and transfection were performed as described in the previous paper [16]. Briefly, human embryonic kidney (HEK) 293A (Thermo Fisher Scientific) cells were cultured in Dulbecco’s Modified Eagle Medium (DMEM, Nissui Pharmaceutical) with 5% fetal bovine serum (FBS, GIBCO, Thermo Fisher Scientific), 6 mg/L penicillin (SIGMA), 10 mg/L streptomycin (GIBCO) and 200 µM glutamine (GIBCO) added (complete DMEM). β-arrestin1/2-deficient HEK293A cells (ΔARRB1/2-HEK293A) derived from HEK293A cells were cultured in complete DMEM with 10% FBS. Cells were cultured in a humidified 37°C incubator containing 5% CO2.

### Transfection

For microscopic observations, transfections were conducted by using Lipofectamine 3000 (Thermo Fisher Scientific) as described in the previous paper [1]. Two coverslips (Matsunami, diameter: 25 mm, thickness: 0.13∼0.17 mm) were placed in a 60 mm dish (Greinar, medium volume: 4 mL) and 2 x 10^6^ cells were seeded and transfected on the same day. Solution A [60 µL of Opti-MEM with totally 1 µg of plasmid (The amount of plasmid was standardized to 1 µg using pCAGGS vector) and 2 µL p3000 reagent] and solution B [60 µL of Opti-MEM with 2.5 µL of Lipofectamine 3000 reagent] were mixed and incubated for 15 minutes at room temperature. The mixture was added to cell culture dishes.

For flow cytometry and the assays using plate readers, transfections were conducted by using PEI MAX (Polysciences). When using 60 mm dishes (Greiner, medium volume: 4 mL), 1.2 x 10^6^ cells were seeded the day before transfection. The amount of plasmid was standardized to 2 µg using pCAGGS (empty vector). Solution A [200 µL of a mixture of plasmid and Opti-MEM (Thermo Fisher Scientific)] and solution B [200 µL mixture of PEI and Opti-MEM at a ratio of 1: 19 were prepared] were mixed and incubated for 20 minutes at the room temperature. Then, the mixture was dropped onto cell dishes. When plates other than 60 mm dish were used, the number of cells and the amount of plasmid were prepared in proportion to the basal area.

### Multicolor single-molecule imaging system

Single-molecule imaging was performed as described in the previous paper [1], but with some instrument upgrades (Figure 1, Supplementary Table 1). Five lasers (OBIS 405-nm, 488-nm, 561-nm, 637-nm lasers, Coherent, and a 703-nm fiber laser, MPB Communications Inc.) were integrated into a single optical axis using four dichroic filters (Di02-R405-25-D, Semrock, RT532rdc-UF2, Chroma, LM01-613-25, Semrock, LM01-659-25, Chroma) in the ascending order of wavelength. For lasers with wavelengths of 488 nm, 637 nm, and 703 nm, excitation filters (FF01-480/17-25, Semrock, ZET642/20x, Chroma, and FF01-716/40-25, Semrock) were placed in front of each laser to cut off excitation light in unexpected wavelength bands. The electric shutters (SSH-25RA with SSH-C2B, OptoSigma) were placed in front of each laser to individually control on and off. Two achromatic lenses (DLB-20-40PM and DLB-30-150PM, OptoSigma) were placed in the optical axis to expand and collimate the lasers. For the 703-nm laser, an additional beam expander with diopter movement (LBED3, OptoSigma) was placed in front of the laser to correct a chromatic aberration. In addition, an achromatic 1/4-wave plate was inserted in the optical axis to convert the linear polarization of the laser to circular polarization. The integrated and collimated laser beams were projected into a fluorescent illuminator (Ti2-LA-BF fixed main branch, Nikon) of the microscope (Ti2E, Nikon) via a two-axis galvanometer scanner. The positions of the XY-galvanometers and the three achromatic lenses (Lens 1: DLB-30-80-PM, Lens 2: DLB-30-100PM, Lens 3: DLB-30-80-PM) were adjusted so that the position of the galvanometer scanners was conjugate to the image plane (Figure 1). Fluorophores on a coverslip were excited by TIRF illumination through a five-band dichroic (ZT405/488/561/640/705rpc, Chroma) and an objective lens (CFI Apochromat TIRF 100XC Oil, Nikon). The laser angle was electrically controlled by the XY-galvanometers through a function generator (AFG-3022, TEXIO). To achieve uniform excitation of the fluorophores in the field of view, rotating laser illumination was typically used as previously reported (Yanagawa, Sako 2021). The fluorescence images were detected by two sCMOS cameras (ORCA Fusion-BT, Hamamatsu) after being split into four color channels using image splitting optics (a W-view Gemini-2C with two W-view Geminis, Hamamatsu). An autofocus system (PAF, ZIDO) was used to keep the focus on the plasma membrane. The entire TIRF microscope system was controlled by the AIS (ZIDO).

**Figure 1.**
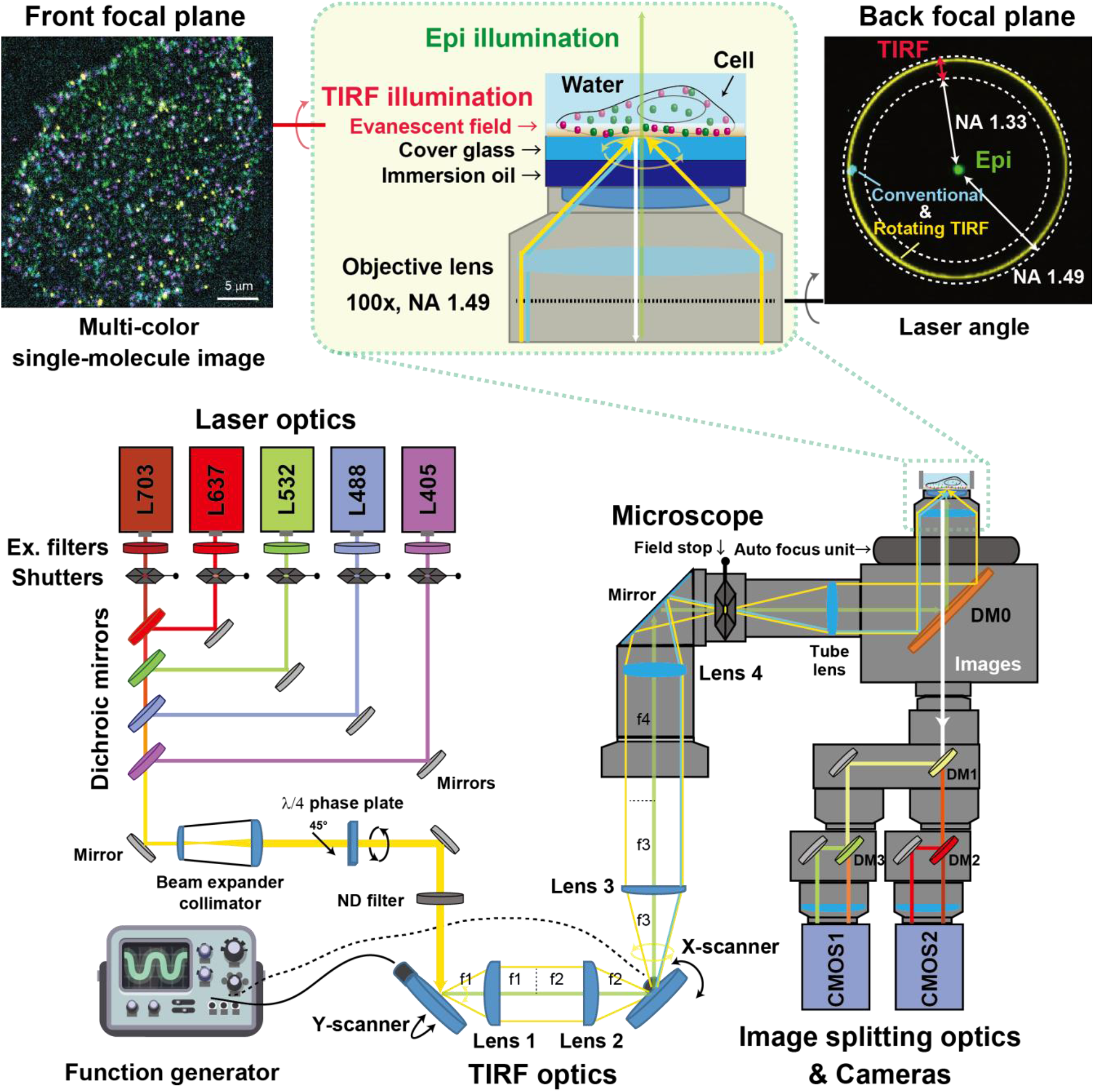
Schematics of multicolor single-molecule imaging system. Detailed configurations are provided in Supplementary Table S1.

### Single-molecule imaging

Transfection was performed one day before observation. For vasopressin type 2 receptor (V2R) observation in four colors, 100 ng of Lg-V2R/pFC15A was transfected. For angiotensin II type 1 receptor (AT1R) and CLC observation in two colors, 100 ng of AT1R-Halo/pFC15A or Lg-AT1R/pFC15A, and 50 ng of EGFP-CLC/pEGFP-C1 were transfected. After overnight incubation, Lg-V2R was simultaneously labeled with 20 nM ST488-FiBiT, 40 nM TMR-FiBiT, 20 nM SF650-FiBiT, 5 nM SF720-FiBiT. AT1R-Halo was labeled with 30 nM HaloTag SF650 ligand (GORYO CHEMICAL). Lg-AT1R was labeled with 10 nM SF650-FiBiT. Phenol red-free DMEM (FluorBright DMEM, Gibco) supplemented with 5% FBS, 6 mg/L penicillin, 10 mg/L streptomycin and 200 µM glutamine (complete phenol red-free DMEM) was used to dilute the fluorescent dyes and F-FiBiT. In each case, 3 mL of complete phenol red-free DMEM containing the fluorescent dye was placed in a 60 mm dish and incubated at 37°C for 15 minutes. After incubation, the medium was changed three times with 3 mL phenol red-free DMEM, and the coverslip was mounted on a chamber (Attofluor Cell, Thrmo Fisher Scientific) for observation. The medium was replaced with 450 µL of HBSS containing 0.01% BSA (BSA-HBSS) before observation. For observation, we visually selected the cells that have bright spot density appropriate for single-molecule tracking (SMT) analysis. Cells with a plasma membrane density greater than 1 µm^-1^ are not suitable for SMT analysis due to the inability to resolve individual bright spots. For simultaneous four-color imaging of Lg-V2R, 300-frame videos were acquired with an exposure time of 30 ms (pixel size: 65 nm/pix, 512×512 pix). For simultaneous two-color imaging of AT1R and CLC, 100-frame videos were acquired before and 5 min after AngII stimulation with an exposure time of 30 ms (pixel size: 65 nm/pix, 512×512 pix). For non-wash imaging, Lg-AT1R-expressing HEK293 cells were incubated in 1 nM SF650-FiBiT in BSA-HBSS for 30 min on the microscope. Then, the 100-frame videos were repeatedly recorded at the same cell positions at 5-min intervals for 60 min (12 time points).

### Flowcytometry

Cell culture and transfection were performed by using a 10 cm dish (Griner, medium volume: 10 mL). HEK293A cells were transfected with 500 ng Lg-AT1R. After incubation for 1 day, cells were harvested using 1 mL of PBS containing 0.5 mM EDTA followed by 1 mL of BSA-HBSS and seeded at 200 µL each onto 96-well v-bottom-plates (Grainer). Then, the cells were stimulated by AngII in BSA-HBSS for 30 minutes in a humidified 37 °C incubator with 5% CO_2_. Then, the medium was exchanged to 100 nM SF650-FiBiT diluted in BSA-HBSS, incubated on ice for 15 min to prevent receptor turnover. The medium was changed to 100 µL of BSA-HBSS, suspended and filtered through a 40 µm filter prior to measurement. Measurements were performed using a flow cytometer (CytoflexS, Beckman Coulter) with an excitation wavelength of 638 nm and a detection wavelength of 660 nm. For the F-FiBiT saturation binding and on-time measurements, the labeling process was performed at 37°C.

### Bioluminescence internalization assay

ΔARRB1/2 - HEK293A cells were transfected with 2 ng of Lg-V2R/pFC15A or Hi-V2R, 200 ng of β-arrestin2 or pCAGGS as a negative control. After incubation for one day, cells were harvested in 2 mL of 0.01% BSA-HBSS, seeded in a 96-well F-bottom white plate (655904, Griner) at a volume of 45 μl per well. A mixture of luminescent substrate (Furimazine 50 µM) and peptide (LgBiT or TMR-HiBiT or TMR-FiBiT 0.4 µM in 0.01% BSA-HBSS) was added in 50 µL per well. After incubation at room temperature for 40 min, baseline luminescence was measured using a luminescence plate-reader (SpectraMaxL, Molecular Devices). 5 µL of arginine vasopressin (AVP) diluted in 0.01% BSA-HBSS was added in each well and luminescence was measured at 30 s interval for 30 min.

## Data analysis

Data analysis for single-molecule imaging was performed based on the following paper [17]. First, the acquired image data were background subtracted using the imageJ Rolling-ball-subtraction algorithm with a ball radius of 25 pixel. To correct for misalignment between channels, affine transformation was performed based on an image of fluorescent beads for calibration (TetraSpeck Microspheres, Invitrogen) using AAS 2.8 (Zido). Then, the SMT and Variational Bayesian and Hidden Markov models (VB-HMM) analyses were performed to estimate the trajectories of particles and assign diffusion states with different diffusion coefficients to each step of the trajectory [26]. The VB-HMM analysis was based on a model which assumes the number of diffusion states and a Markov chain between each state. In this study, we compared the variational lower bounds computed based on the estimation results of the 1-5 state models. The lower bound is calculated based on the explanatory power of the estimation results (the residual between the time-series change in trajectory step size and the estimation results) and the Kullback-Leibler information content (a penalty term based on the prior distribution and number of parameters). In the present study, a four-state model was selected that maximized the lower variational bound in many cells as previously reported [14,15]. Co-localization between two molecules was defined by the two bright spots being within 100 nm of each other in the same frame and classified in the same diffusion state.

The calculation of the parameters from trajectory and the curve fitting were obtained with Igor Pro9.02 (Igor, WaveMetrix) as follow. The MSD within time nΔt of each trajectory was calculated by

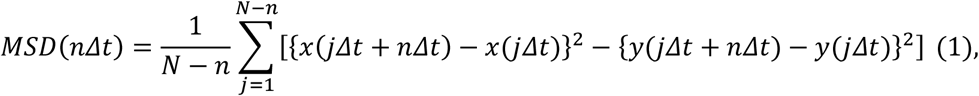

where n is the length of frames, Δt is the frame rate, and N is the total frame number of the trajectory. The MSD-Δt plots are fitted using the following equation.

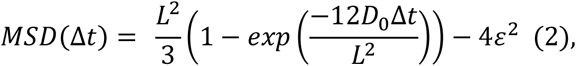

where L is the confinement length, D_0_ is the diffusion coefficient taking the limit of Δt to 0, and ε is an error term.

Flow cytometry analysis was performed using FlowJo 10.10.0. After gating live cells with forward and side scattered light, histograms were generated, and Mean-Fluorescence-Intensity (MFI) was calculated.

For internalization detection experiments using a luminescence plate reader, data were analyzed using Excel. The mean luminescence values measured before agonist stimulation were corrected to 1. To correct for luminescence decay during the measurement, the mean intensity of each well was normalized to the intensity during vehicle (without ligand) stimulation.

The concentration-response curves with logarithmic x-values were fitted with

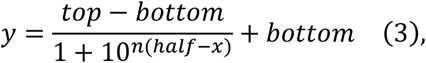

where n is the Hill-slope, half is the log_10_(EC_50_) and (top - bottom) is the span (E_max_). A four-parameter curvilinear regression model was adopted and fitted using prism10 with the absolute value of the Hill-slope within 1.5.

One site binding curve in saturation binding was fitted with

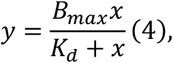

where B_max_ is the maximum binding, K_d_ is equilibrium dissociation constant. Graph s and statistical tests were performed using PRISM 10 (Graphpad) or Igor. Unless otherwise noted, error bars represent SEM.

## Results

### Development of a four-color single-molecule imaging system

To simultaneously monitor multiple molecules within a single cell, we developed a four-color TIRF imaging system (Figure 1 and Supplementary Table 1). The five collimated lasers (405, 488, 561, 637 and 703 nm) are integrated into one optical axis and projected into a microscope. The XY scanning galvo mirrors are placed at the conjugate position to the image plane and the angles of the lasers are electrically controlled by the function generator. To achieve uniform TIRF illumination, the 90° phase-shifted 300 Hz sine waves are sent to the XY-galvo drivers to produce a rotating beam at the back focal plane of the objective within NA 1.33-1.49. By adjusting the amplitude of the voltage in each of the XY galvo drivers, the lasers were adjusted to focus on a circle around the circumference of the aperture. The position of the laser in the back focal plane of the objective was checked through the Bertrand lens. Fluorescence images were spectrally split into four wavelength bands and captured simultaneously by two sCMOS cameras.

### Fluorescent FiBiT peptide-based GPCR labeling for multicolor single-molecule imaging

Taking advantage of the high affinity binding between HiBiT peptide and LgBiT fragment, we developed a specific fluorescence labeling method for simultaneous four-color single-molecule measurements. As a first attempt, we tried to label Lg-GPCRs with TMR-HiBiT peptide (TMR-HiBiT, Figure 2A,B, and C, left panel). However, the addition of 10 nM TMR-HiBiT resulted in strong non-specific binding to the coverslips even after extensive wash, probably due to its hydrophobic nature, making it difficult to use TMR-HiBiT for single-molecule imaging. To reduce the non-specific binding, we next synthesized a TMR-FLAG-HiBiT peptide (TMR-FiBiT). The fusion of FLAG-tag, a commonly used hydrophilic tag, greatly reduced the non-specific binding of TMR-FiBiT on coverslips compared to TMR-HiBiT (Figure 2C, right panel). We then additionally synthesized ST488-, SF650-, SF720-FiBiT peptides to generate color variants of fluorescent FiBiT (F-FiBiT). In particular, the SF650-FiBiT is ideal in terms of photostability and non-specific binding among the four F-FiBiTs.

**Figure 2.**
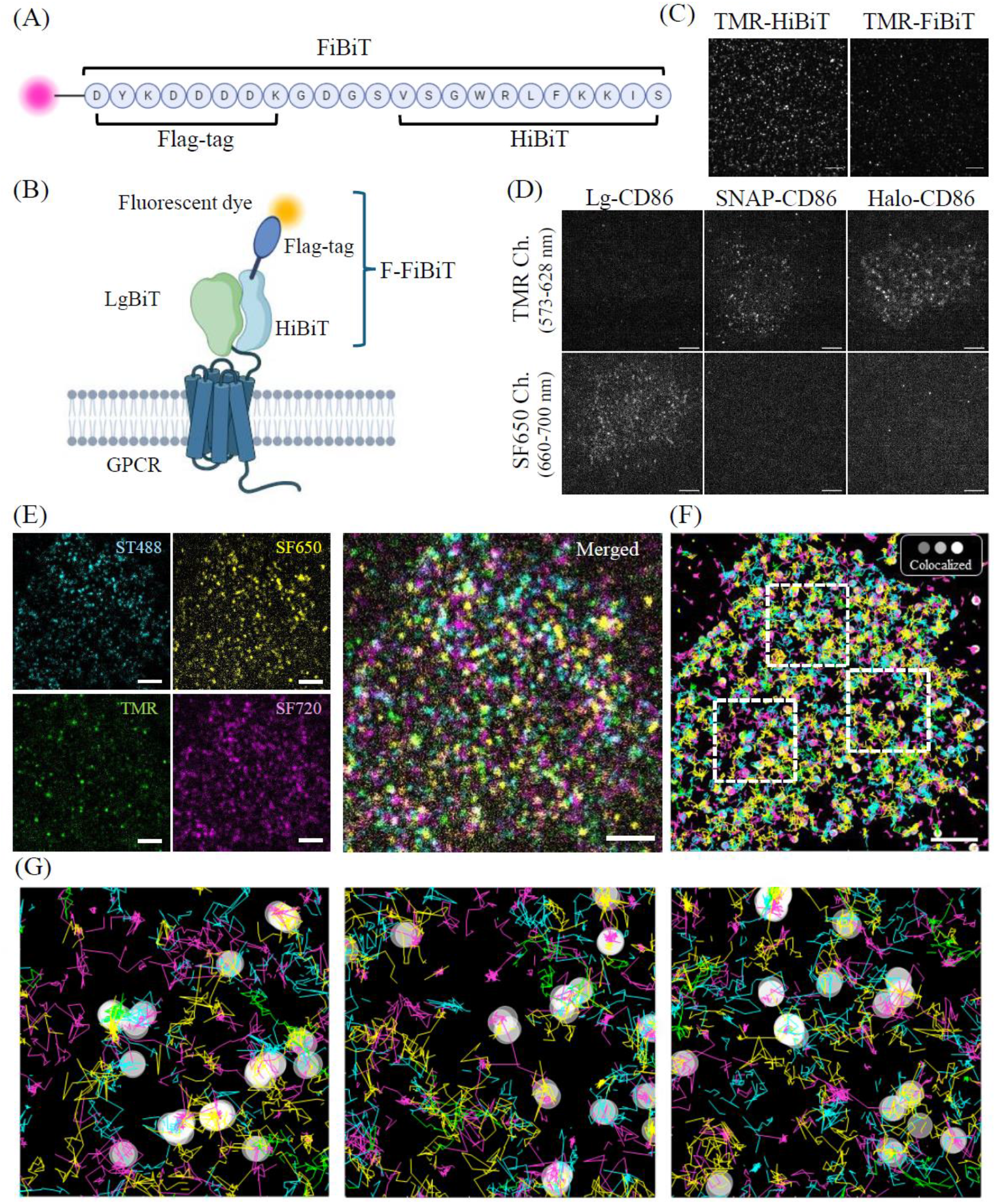
(A) FiBiT peptide sequence. (B) Schematic of GPCR labelling using HiBiT. (C) Non-specific binding of TMR-HiBiT (left) and TMR-FiBiT (right) on coverslips. The 10 nM TMR-HiBiT or TMR-FiBiT was added to a 60 mm dish with a coverslip and washed three times with complete phenol red-free DMEM, then the medium was replaced with BSA-HBSS for observation. (D) Two-color single-molecule images of Lg-CD86 (left), SNAP-CD86 (middle), Halo-CD86 (right) expressing HEK293 cells simultaneously labeled with 10 nM SF650-FiBiT, 10 nM SNAP-TMR, and 10 nM Halo-JF549 mixture solution. Representative images of TMR/JF549 and SF650 channels are shown in the upper and lower panels, respectively. (E) Example four-color single-molecule image of Lg-V2R in the basal state (no ligand). HEK293A cells expressing Lg-V2R were simultaneously labeled with 20 nM ST488-FiBiT, 40 nM TMR-FiBiT, 20 nM SF650-FiBiT, 5 nM SF720-FiBiT (left), merged images of four channels (right). (F) Trajectories of F-FiBiT-labeled Lg-V2R in (E) were projected on the colocalized coordinates (round markers). Markers were displayed brighter according to the degree of co-localization. (G) shows enlarged views of the area enclosed by the white dotted lines (5 µm squares). Mean duration of colocalization events is ∼50 ms in the cell shown in (E). Scale bars: 5 µm (C, D), 3 µm (E, F).

Next, we evaluated whether the F-FiBiT labeling method could be used simultaneously with the conventional labeling methods using SNAP-tag and HaloTag. We used the CD86 (1-277) as a control monomeric transmembrane protein. LgBiT-, HaloTag-, or SNAP-tag-fused CD86 (Lg-CD86, Halo-CD86, SNAP-CD86) expressing HEK293A cells were simultaneously labeled with SF650-FiBiT, SNAP-cell TMR-star ligand (SNAP-TMR), and HaloTag Janeria Fluor 549 ligand (Halo-JF549). The Lg-CD86-expressing HEK293 cell was exclusively stained by SF650-FiBiT but not by SNAP-TMR nor Halo-JF549. Conversely, the SNAP-CD86 and Halo-CD86-expressing cells were labeled by SNAP-TMR ligand and Halo-JF549, respectively, but not by SF650-FiBiT (Figure 2D). These results indicate that the F-FiBiT labeling method can be used with the SNAP-tag and HaloTag labeling method.

We examined feasibility of F-FiBiT labeling for multicolor single-molecule imaging. Lg-V2R expressing HEK293 cells were simultaneously stained by the four-color F-FiBiT peptides (Figure 2E). Lg-V2R molecules in the same plasma membrane were simultaneously monitored in four different channels with an exposure time of 30 ms (Figure 2E). The single-Lg-V2R molecules are transiently colocalized with each other as previously reported for other GPCR (Movie 1, Figure 2F) [7].

### Bioluminescence-based plate reader assay to monitor ligand-induced internalization of Lg-GPCR

To confirm that the N-terminal F-FiBiT-LgBiT labeling on GPCRs does not inhibit their function, we assessed ligand-induced endocytosis of V2R. In the previous studies using the HiBiT system, currently commercially available as Nano Glo HiBiT Extracellular Detection System (Promega), purified LgBiT protein is added to detect cell surface expression of the N-terminal HiBiT-fused GPCR (Hi-GPCR) by bioluminescence from the reconstituted NanoLuc [16,21]. We here compared the conventional LgBiT/Hi-GPCR method and F-FiBiT/Lg-GPCR methods (Figure 3). Using the Lg-V2R and F-FiBiT labeling, bioluminescence signal was detected before agonist stimulation as well as by the conventional method (Figure 3A, B), suggesting that the fusion of the dye and the FLAG-tag at the N-terminal of HiBiT peptide does not affect its function as a split luciferase. To observe β-arrestin-dependent endocytosis of Lg-V2R, we compared the βarr1/2-deficient HEK293A cell line with and without βarr2 over-expression. Consistent with the conventional method, agonist-induced internalization was detected only in cells co-expressing βarr2 (Figure 3C-E, EC_50_ ∼15 nM in both methods). These results suggest that the F-FiBiT/LgBiT fusion does not inhibit the ligand-induced β-arrestin recruitment and subsequent endocytosis of V2R.

**Figure 3.**
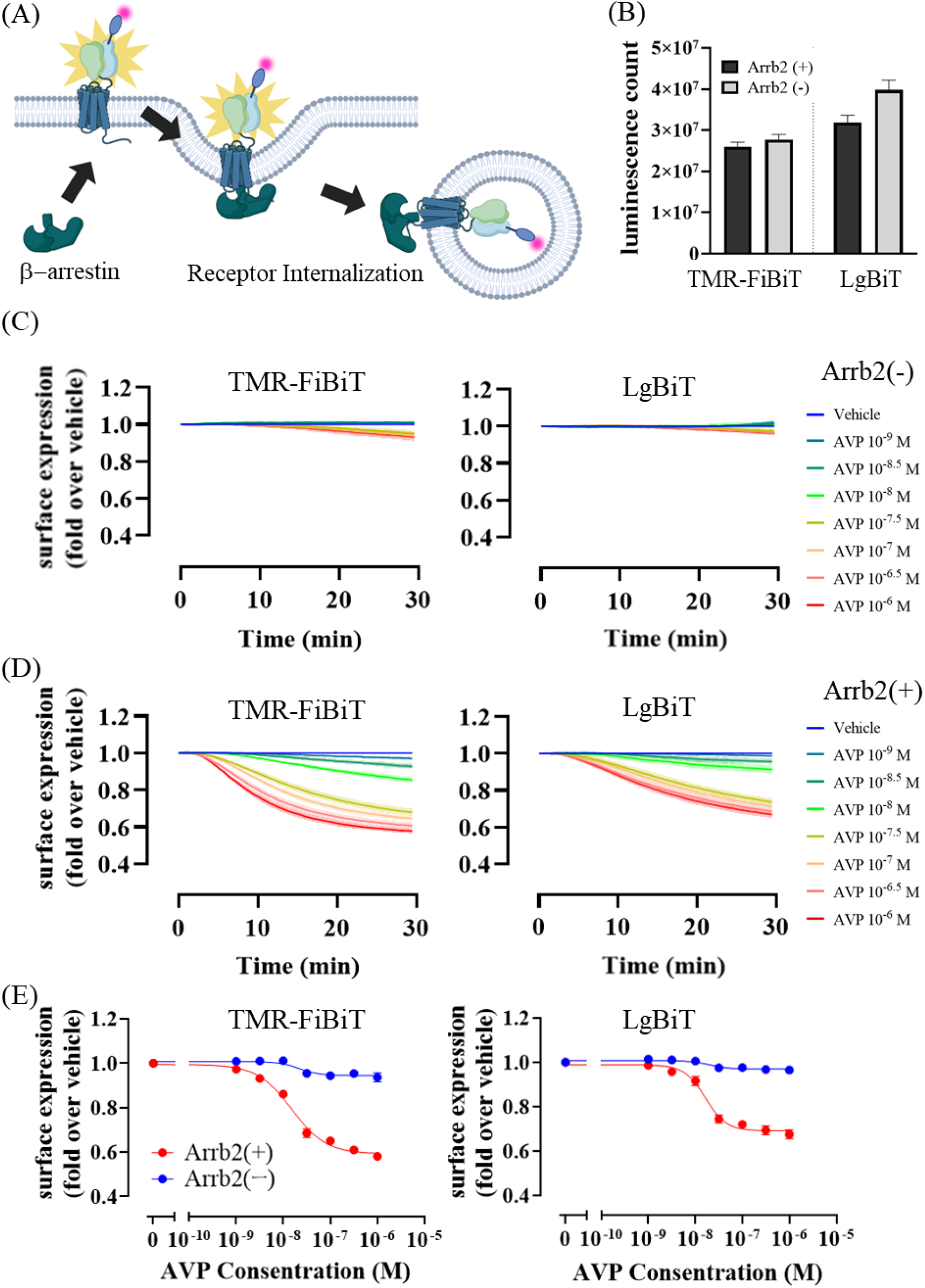
(A) Schematic of a plate reader-based internalization assay to detect luciferase luminescence values. (B) Luminescence count before agonist stimulation. (C, D) Time-dependent changes in luminescence counts and (E) AVP concentration-response-curve in luminescence counts under conditions where Lg-V2R was expressed and TMR-Flag-HiBiT was added (left), HiBiT-V2R was expressed and LgBiT was added (right). The luminescence counts of the condition with BSA-HBSS was corrected to 1. (C) Shading and (D) error bars represent SEM (n=3).

### Flow cytometry-based estimation of affinity between F-FiBiT and Lg-GPCR on living cell surface

To assess the dissociation constant (K_d_) between Lg-AT1R and SF650-FiBiT, we measured the labeling rate of Lg-GPCRs by F-FiBiT in an AngII concentration-dependent manner (Figure 4A). We compared the saturation binding of SF650-FiBiT on Lg-AT1R-expressing and mock-transfected HEK293 cells, and specific binding to cell surface Lg-AT1R was estimated as differential mean fluorescence intensity (MFI) using flow cytometry. The K_d_ was estimated to be ∼83 nM based on the one-site binding model (Figure 4A), which means that 10-20% of Lg-GPCRs were labeled at the concentrations observed in the single-molecule imaging (10 nM). We also confirmed that F-FiBiT-labeled GPCRs were able to observe for at least 1 hour without significant decay, indicating the low off-rate (Figure 4B).

**Figure 4.**
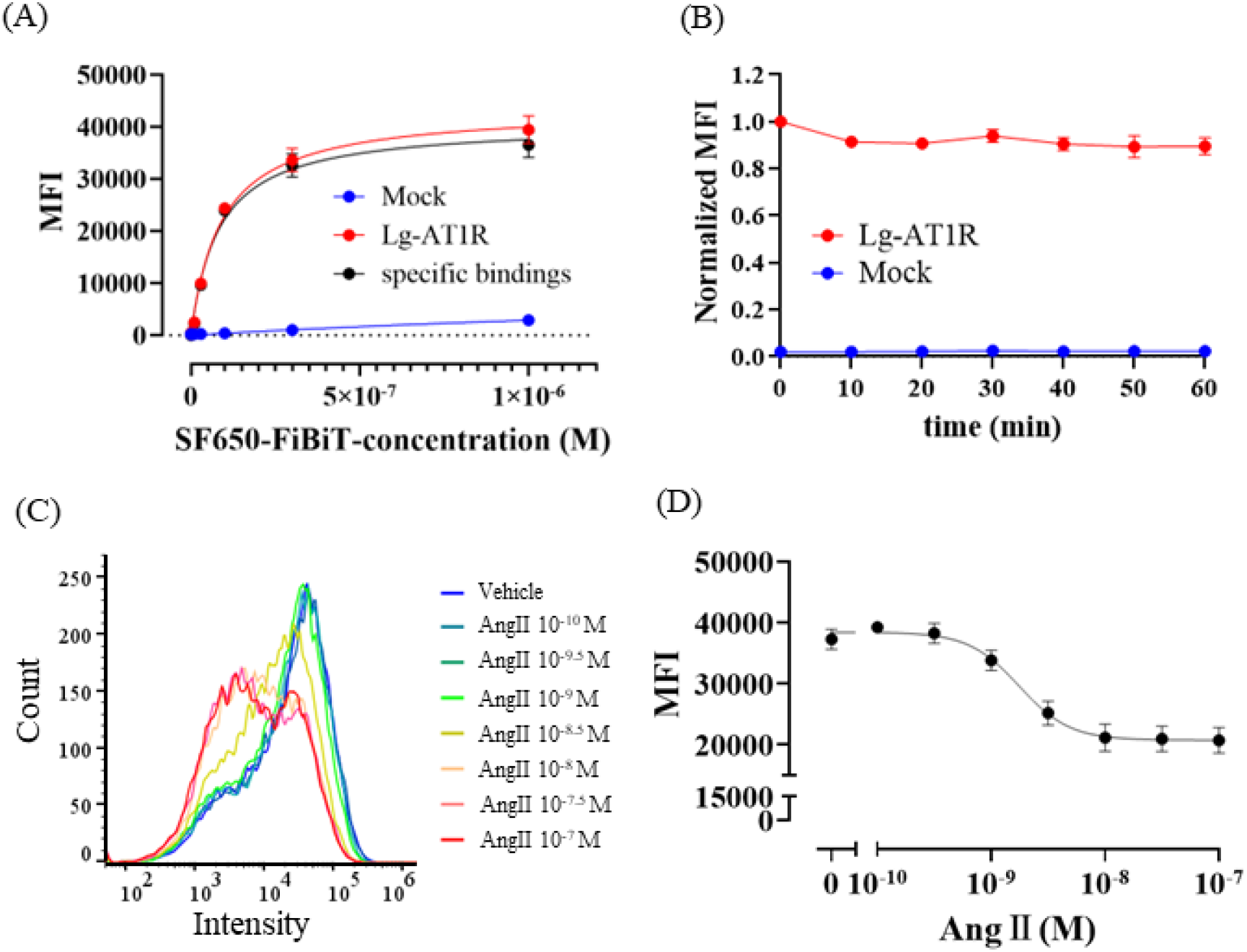
(A) Concentration-dependent binding of SF650-FiBiT to Lg-AT1R expressing (Lg-AT1R) or non-expressing (Mock) HEK293A cells. Specific binding was calculated as the difference in mean fluorescence intensity (MFI) between Lg-AT1R- and Mock-transfected cells. (B) Time-dependent changes in MFI of SF650-FiBiT-labeled Lg-AT1R in HEK293 cells after washing (red). Non-specific binding was estimated from mock-transfected HEK293A cells. Lg-AT1R- and Mock-transfected cells were labeled with 100 nM SF650-FiBiT. Data were normalized to MFI of Lg-AT1R expressing cells at 0 min. (C) Representative intensity histograms of Lg-AT1R-expressing HEK293 cells 30 min after Ang II stimulation with different concentrations. (D) Ang II concentration dependent change in MFI of Lg-AT1R-expressing HEK293A cells were stimulated with Ang II and then labeled with 100 nM SF650-FiBiT for membrane surface receptors and MFI was measured at 660 nm by flow cytometry. Experiments were performed in duplicate, n=3. Error bars represent SEM.

To test further applications of F-FiBiT in a flow cytometry-based endocytosis assay at the single cell level, we measured the specific binding of SF650-FiBiT to cell surface Lg-AT1Rs 15 min after stimulation by Ang II. The Ang II-induced decrease in the cell surface Lg-AT1Rs was successfully monitored, reflecting the internalization of Lg-AT1R from the cell surface in a concentration-dependent manner (Figure 4C, D). The EC_50_ (∼1.8 nM) was consistent with our previous report using the split luciferase assay [16].

### Single-molecule diffusion dynamics comparison between F-FiBiT and HaloTag labeling methods

To confirm whether GPCRs labeled with the F-FiBiT show similar changes in diffusion dynamics reflecting the recruitment into the CCP as the conventional HaloTag method, a comparative two-color single-molecule imaging analysis was performed using AT1R as another model GPCR. AT1R-Halo or Lg-AT1R were co-expressed with EGFP-CLC in HEK293A cells and labelled with HaloTag SF650T ligand and SF650-FiBiT respectively (Figure 5 A-C). In accordance with the previous studies [16,17], trajectories of AT1R were classified into four states (immobile, slow, medium and fast) based on the time-series of the displacement within 30 ms by the VB-HMM analysis [17,26]. The MSD-Δt plots show that the immobile and slow states are in a confined diffusion mode, whereas the medium and fast states showed a Brownian motion on average (Supplementary Figure S1). In both labeling methods, Ang II stimulation increased AT1Rs in the immobile state and decreased AT1Rs in the medium and fast states (Figure 5D). The diffusion coefficient of each state was also decreased by Ang II stimulation (Supplementary Figure S1). Colocalization analysis showed that most of the AT1R colocalized with CLC were in the immobile state, which significantly increased upon Ang II stimulation (Figure 6A). Ang II stimulation increased both on-rate and on-time of colocalization in either of the labeling methods (Figure 6B,C). Collectively, we were able to observe GPCR diffusion dynamics using both the F-FiBiT labeling method and the HaloTag method.

**Figure 5.**
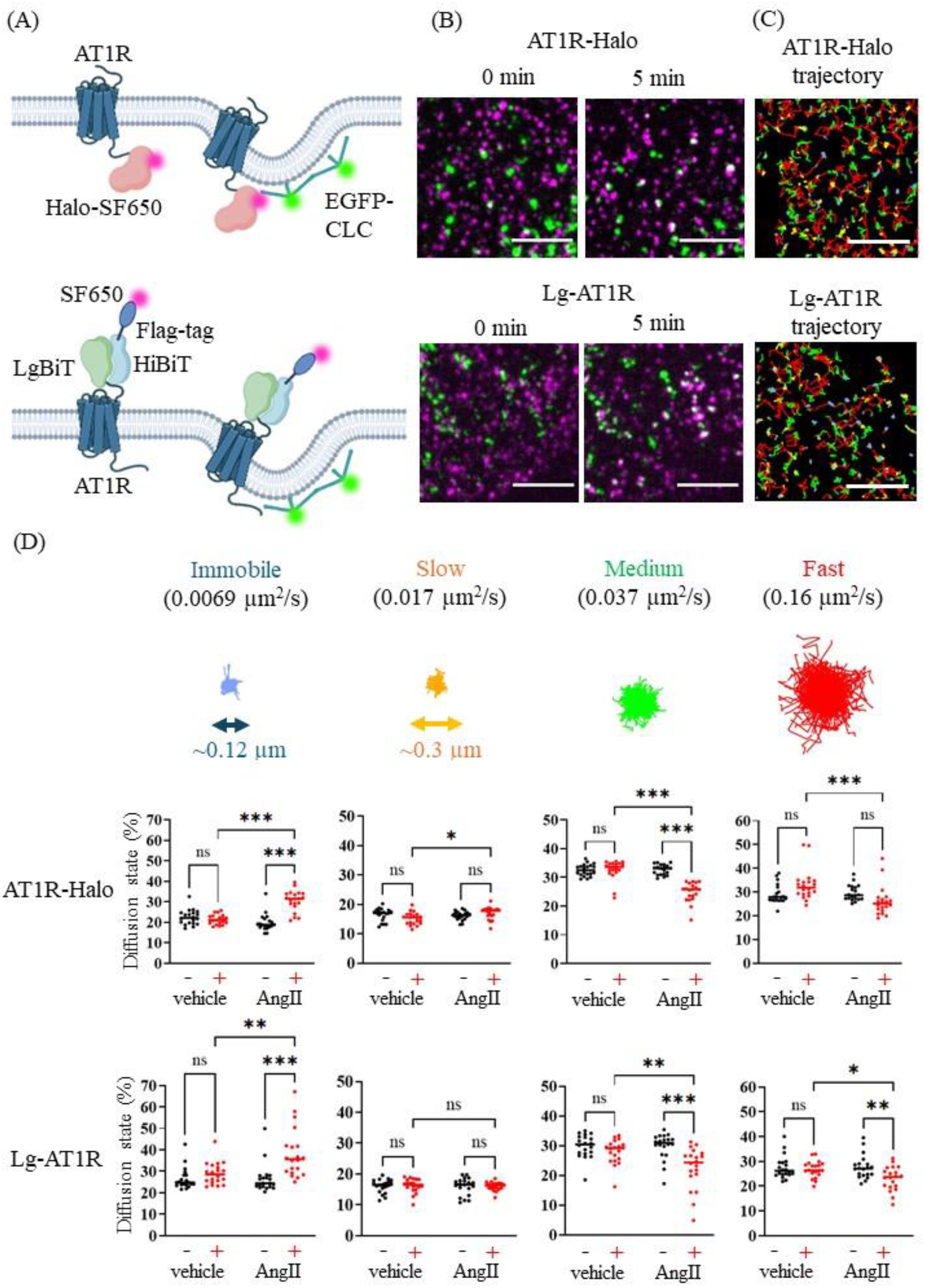
(A) Schematic diagrams of labelling method using HaloTag (top) and F-FiBiT (bottom). (B) Images before (0 min) and after (5 min) Ang II stimulation. Magenta: SF650, green: GFP, scale bar: 5 µm. (C) representative trajectories of four diffusion states of 10 nM AT1R (Blue: immobile, yellow: slow, green: medium, red: fast, scale bar: 3 µm). (D) The percentage of each diffusion state before [0 min (-)] and after [5 min (+)] stimulation. n = 20. (*p < 0.033, **p < 0.002, ***p < 0.001. Two-way ANOVA followed by Tukey’s multiple comparison test)

**Figure 6.**
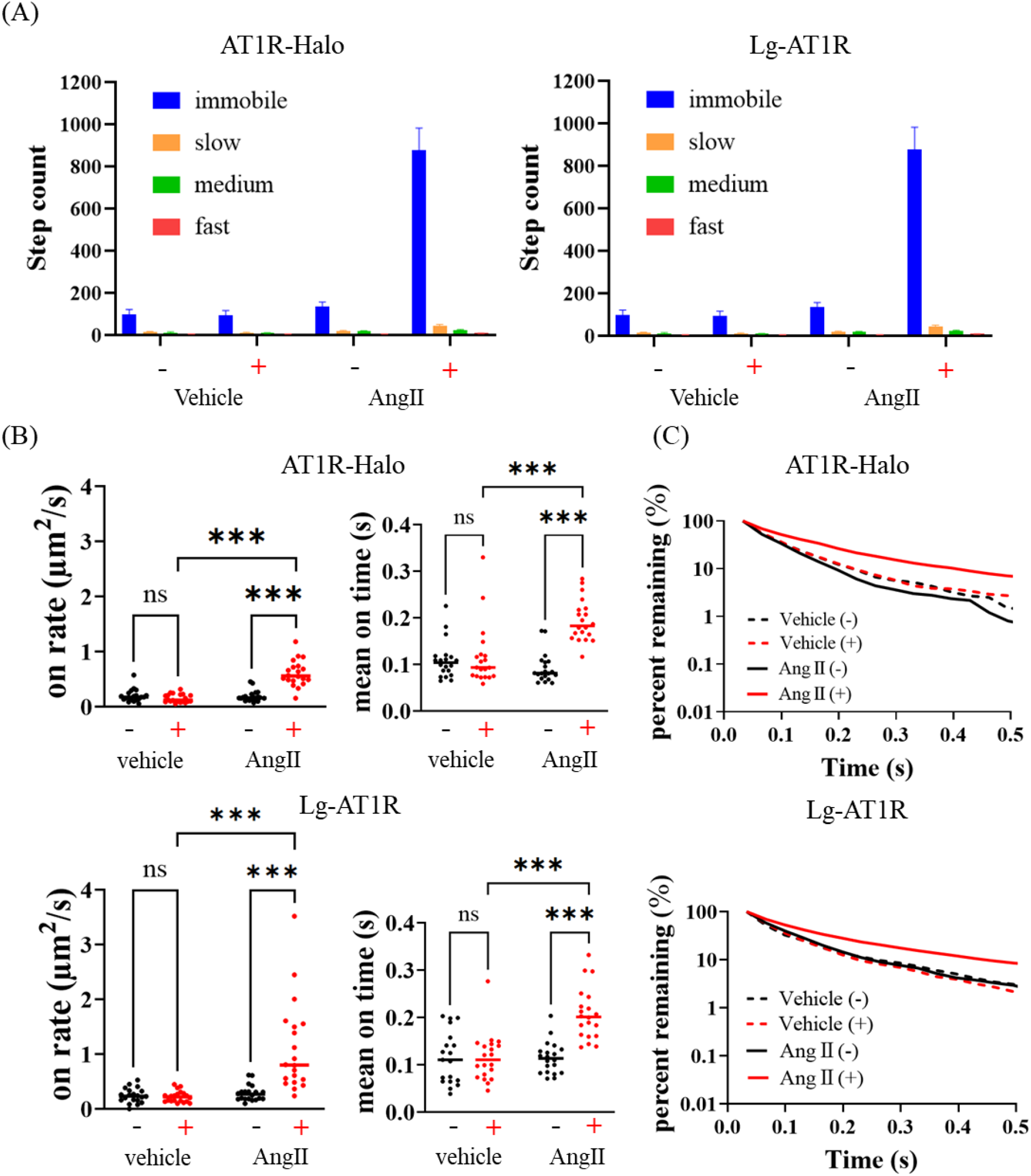
(A) Number of AT1R-clathrin colocalization steps for each diffusion state before (-) and after (+) 10 nM AngII stimulation (left panel: AT1R-Halo, right panel: LgBiT-AT1R). Mean ± SEM (n = 20). (B) On-rate (left panel) and mean co-localization duration (right panel) before [0 min (-) ] and after stimulation [5 min (+) ] (upper panels: AT1R-Halo, lower panels: LgBiT-AT1R). (C) AT1R-clathrin colocalization time distribution. (*p < 0.033, **p < 0.002, ***p < 0.001. Two-way ANOVA followed by Tukey’s multiple comparison test, n = 20)

### Non-wash F-FiBiT labeling for long-term timelapse single-molecule imaging

A simplification of the time-consuming staining and washing process is required for the application of F-FBiT labeling method to drug evaluation using an automated-single-molecule imaging platform [27–30]. We tested the feasibility of a non-wash protocol to label a portion of the cell surface overexpressing Lg-GPCR. After 30 min incubation in 1 nM or lower concentration of SF650-FiBiT/BSA-HBSS, the Lg-AT1R were successfully monitored at the single-molecule level (Figure 7A).

**Figure 7.**
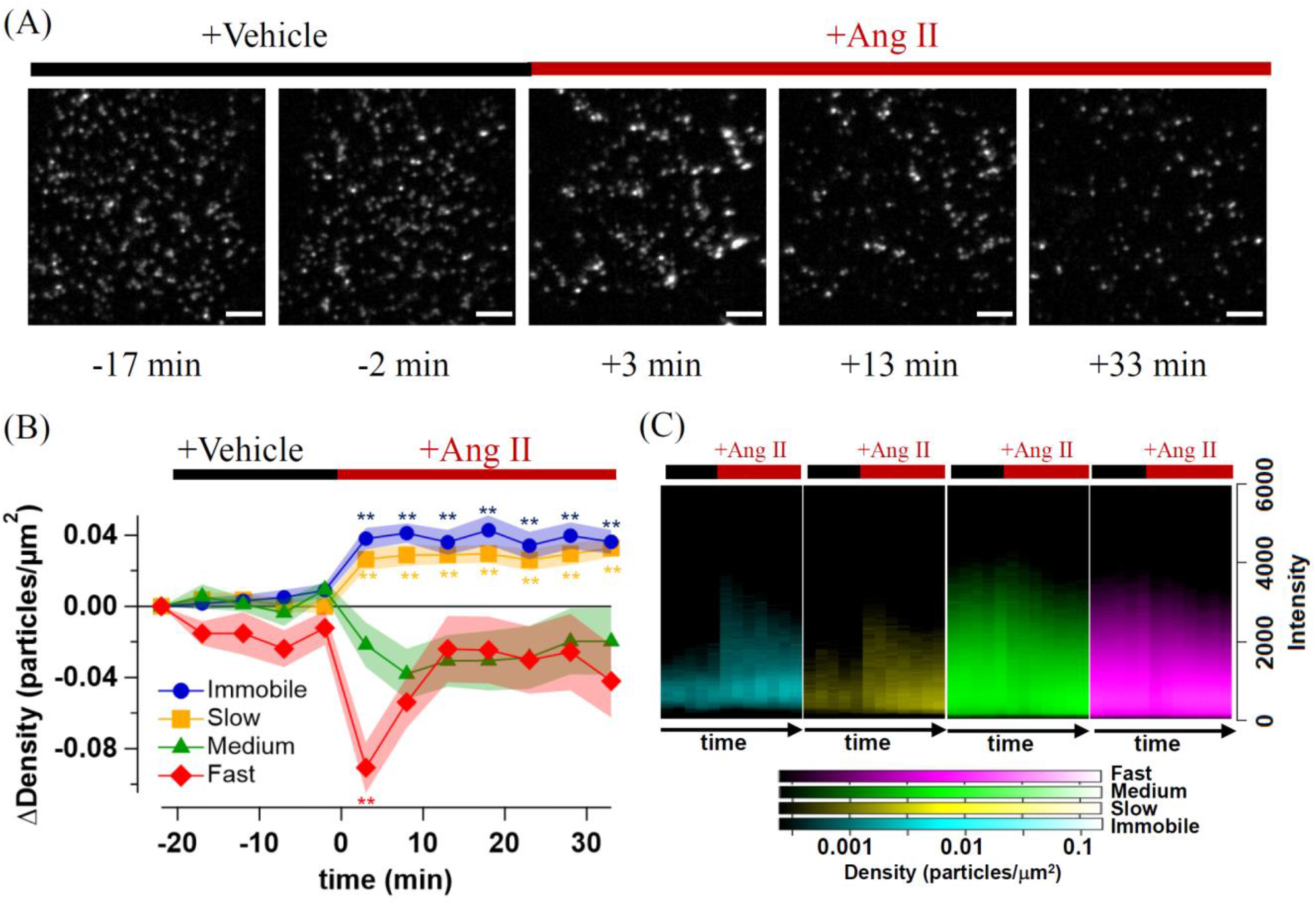
(A) Timelapse images of an identical Lg-AT1R expressing HEK293 cells stained with 1 nM SF650-FiBiT (Scale bar: 3 µm). Time is shown with 1 μM Ang II addition as 0 min. (B) Time-dependent changes of particle density in each diffusion state. Mean ± SEM (n = 18 cells). **p < 0.01 of one-way ANOVA followed by Dunnett’s multiple comparison test versus pre-vehicle stimulation. (C) Heat maps showing the time course of intensity histograms for each diffusion state. The particle density with each intensity was displayed according to a color scale.

The no-wash protocol is expected to be suitable for long timelapse measurements because it allows steady-state measurements with less than 1% of cell surface Lg-GPCR stained by the low concentration of SF650-FiBiT in the outer solution. We took 100-frame videos (3 s) of Lg-AT1R expressing cells every 5 min for 12 time points (Figure 7A). The vehicle and 1 μM Ang II were sequentially added 3 min before time points 2 and 6. SF650-FiBiT/Lg-AT1R diffusion and intensity histograms were not altered by vehicle stimulation (Figure 7B,C). After Ang II stimulation, the particle densities in the immobile and slow states increased significantly and continuously, but that of fast state was transiently decreased 3 min after stimulation (Figure 7B). The brighter particles in the immobile and slow states were increased 3 min after AngII stimulation and gradually decreased over time (Figure 7C), probably due to the accumulation of Lg-AT1R into CCP followed by endocytosis as shown in Figures 3-6.

## Discussion

In the present study, we synthesized the F-FiBiT peptides to expand the use of the HiBiT-LgBiT split luciferase system to cell-surface fluorescent labeling of membrane receptors. There are several advantages for F-FiBiT compared to conventional methods. First, the size of the LgBiT with F-FiBiT (∼19 kDa) is smaller than the conventional tags such as HaloTag (∼33 kDa), and mEGFP (27 kDa), and is comparable to those of SNAP and CLIP-tags (∼19 kDa), reducing a possible artifact on the function of the protein of interest. Second, F-FiBiT is applicable to both fluorescence and bioluminescence measurements. This unique feature allows us to monitor receptor dynamics on the cell surface over a wide range of spatio-temporal scales, from the single-molecule imaging to the high-throughput plate-reader-based endocytosis assays. Third, the high affinity of SF650-FiBiT for LgBiT and its low non-specific binding to coverslips allows for non-washing and non-blocking measurements in single-molecule imaging and flow cytometry, respectively (Figures 4 and 7). This is superior to conventional fluorescent antibody-based endocytosis assays that require blocking, staining, and washing steps, both in terms of time and cost. The non-wash protocol, which avoids cell peeling during solution exchange, is also useful in high-throughput automated single-molecule imaging for drug screening using 96- and 318-well plates [27–30]. Taken together, the F-FiBiT labeling method is a convenient system that only requires a single plasmid DNA to perform multiple assays.

The development of a multicolor single-molecule imaging system, combined with the F-FiBiT labeling method, successfully monitors the Lg-GPCR molecules in the same cell in four colors simultaneously (Figure 2E). This is useful for accurate assessment of GPCR dimerization compared to single-color intensity histogram analysis. Given the diffraction limit of optical microscopy (∼300 nm) is an order of magnitude higher than the localization precision of single molecules (10-20 nm), colocalization analysis based on multicolor single-molecule imaging is advantageous for accurate assessment of GPCR dimerization compared to single-color intensity histogram analysis. This labeling technique may be useful for distinguishing higher-order oligomerization of GPCRs in future studies.

The limitation of F-FiBiT is the membrane impermeability, which prevents us from labeling the LgBiT fusion proteins in the cytoplasm. We have previously discussed the membrane permeability of fluorescent labeling dyes as a factor that may account for the apparent difference in agonist-dependent changes in diffusion dynamics of GPCRs between literatures [17,31]. Constitutive exocytosis and endocytosis of the receptor molecules after labeling could alter the composition of visible receptor molecules at the plasma membrane. In the present study, to examine whether the membrane impermeability makes an apparent difference in the molecular behavior of GPCRs, we compared the changes in the diffusion dynamics of AT1R-Halo stained with the membrane-permeable SF650-HaloTag ligand and Lg-AT1R labeled with F-FiBiT. Contrary to the previous assumption, similar changes in diffusion dynamics and accumulation to the CCP of AT1R-Halo and Lg-AT1R were observed, regardless of whether the intracellular AT1R was labeled or not (Figure 5 and 6). Furthermore, similar changes were observed in the non-wash protocol, where Lg-AT1R on the cell surface could be continuously stained by 1 nM SF650-FiBiT (Figure 7). Therefore, the endocytosis and exocytosis after labeling does not cause a change in receptor behavior upon activation, indicating that the F-FiBiT labeling method is useful for single-molecule diffusion-based estimation of drug effects on membrane receptors. Future development of membrane-permeable F-FiBiT is required to extend the F-FiBiT labeling method to intracellular molecules with LgBiT-tag.

Because F-FiBiT can be used with HaloTag, SNAP-tag, and fluorescent proteins, it may be used to monitor four different molecules simultaneously, each with a different dye. Simultaneous four-color single-molecule imaging can help elucidate spatiotemporal GPCR signaling in the future. For example, it will be an important method to elucidate the dynamics of the “megaplex”, a complex consisting of GPCRs, G proteins, and β-arrestins that may regulate the ERK pathway in living cells [32]. The development of more stable and brighter dyes in order to achieve highly efficient, prolonged multicolor single-molecule imaging is a challenge. Currently, mEGFP, SNAP-TMR, Halo-SF650, and SF720-FiBiT are the better set of tag-dye combinations for four-color imaging, but mStayGold will be the superior gold standard for the GFP channel [33]. In addition, a brighter and hydrophilic dye for the SF720 channel is also needed to improve image quality. By combining F-FiBiT with superior fluorescent dyes and orthogonal labeling methods, we will be able to better understand the dynamics of a diverse supramolecular assembly formed in cells at the single-molecule level.

## Conclusion

In conclusion, we have established a method to label cell surface-expressing LgBiT-GPCRs with F-FiBiT peptide. The versatile use of the F-FiBiT peptide adds an option for future multicolor fluorescence and bioluminescence imaging in GPCR pharmacology and cell biology.

## Author Contributions

M.Y. and A.I. conceptualized and supervised this project. T.Y. and M.Y. designed and performed all experiments, statistical analyses, data visualizations, and figure illustrations. Y.S and M.Y. developed the microscopy system and the single-molecule analysis programs. A.I. developed the bioluminescence plate reader- and flowcytometry-based endocytosis assay protocols. M.Y. and T.Y. drafted and revised the manuscript with contributions from all authors.

## Data Availability

The evidence data generated and/or analyzed during the current study are available from the corresponding author on reasonable request.

## Acknowledgements

We thank Dr. Reiko Ito (Riken CBS) for the synthesis of the F-FiBiT peptides, Dr. Masato Yasui (ZIDO corp.) for the custom development of imaging and analysis software for single-molecule images, and Ms. Hiromi Sato (Riken CPR) for plasmid DNA construction. We thank Mr. Ayaki Saito and all the members of the Inoue Laboratory for critically reading and editing the manuscript. This work was supported by JSPS KAKENHI (19H05647, JP21H04791, JP21H05113, JP21H05037, 24K01982, 24H01266) to Y.S., A.I., and M.Y., by JST FOREST and Moonshot (JPMJFR215T and JPMJMS2023) to A.I., and JST PRESTO (JPMJPR20EF) to M.Y., by AMED BINDS and Moonshot (JP22ama121038 and JP22zf0127007) to A.I., and RIKEN Pioneering Project “Glyco-Lipidologue Initiative” to Y.S.

