## Supplementary Figure S1 for "Four-color single-molecule imaging system for tracking GPCR dynamics with fluorescent HiBiT peptide"

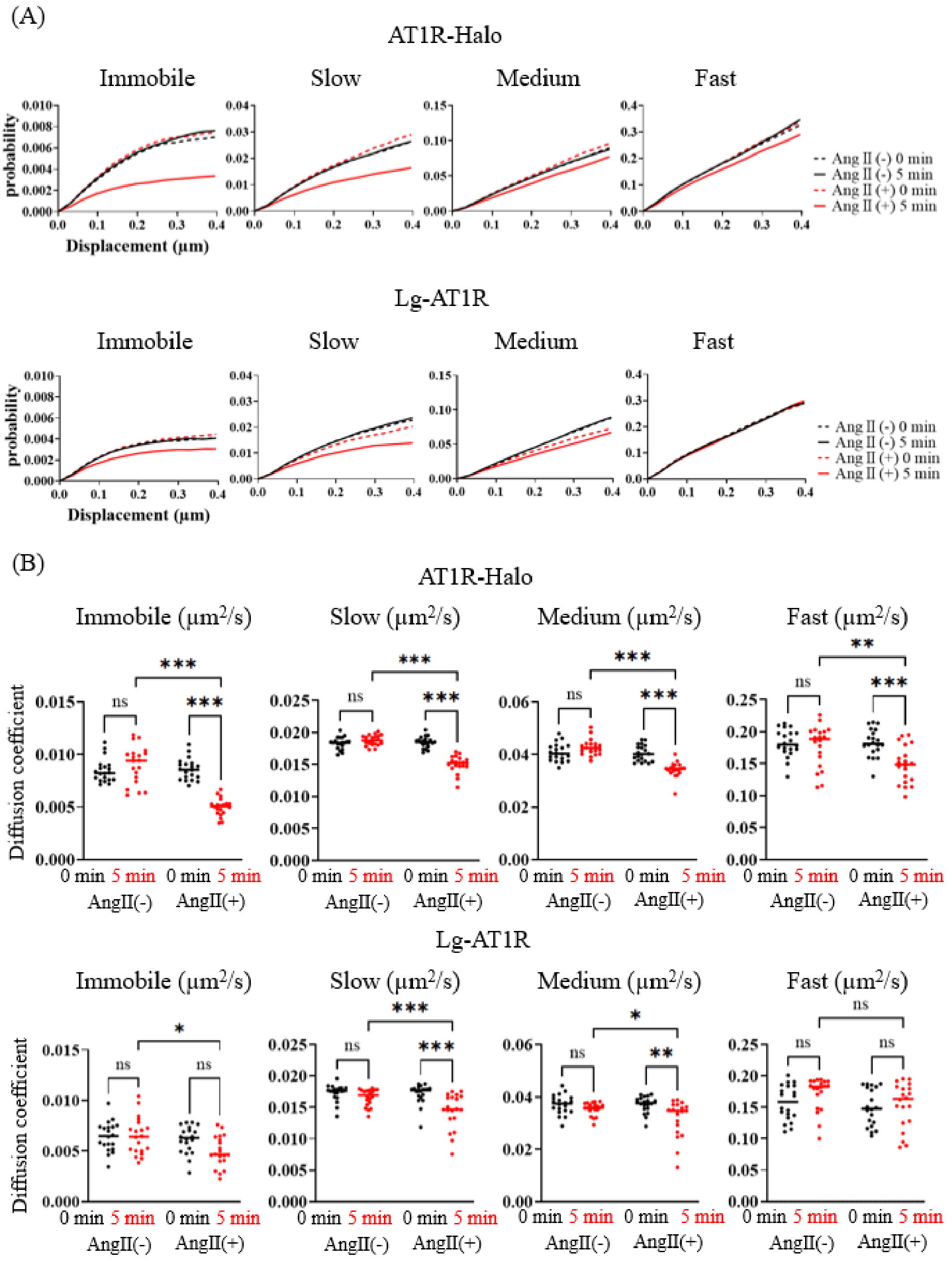


**Supplementary Figure S1**

(A) MSD-Δt plots of SF650-labled AT1R for each diffusion state before [0 min (-) ] and after stimulation [5 min (+) ] (upper panels: AT1R-Halo, lower panels: LgBiT-AT1R). (B) Comparison of the diffusion coefficients of AT1R for each diffusion state before [0 min (-) ] and after stimulation [5 min (+) ]. n = 20. (*$p < 0.033$, **$p < 0.002$, ***$p < 0.001$. Two-way ANOVA followed by Tukey's multiple comparison test)
