## Supplementary Table S1 for "Four-color single-molecule imaging system for tracking GPCR dynamics with fluorescent HiBiT peptide"

**Supplementary Table S1.** List of main configurations of Nikon AIS4C

| No. | Product name | Maker | Catalog number | amount | Use |
|---|---|---|---|---|---|
| 1 | Microscope system | Nikon | Ti2-E FL2-T1 | 1 | microscope |
| 2 | Objective Apo TIRF 100x 1.49 | Nikon | CFI Apochromat TIRF 100XC Oil | 1 | objective lens |
| 3 | Perfect Auto Focus system | ZIDO | PAF | 1 | auto focus |
| 4 | Power supply for PAF, 100 VAC | Thorlabs | GPS011-JP | 1 | auto focus |
| 5 | DA board | Interface | URS-343416 | 1 | auto focus |
| 6 | DA board, USB cable | Interface | ECO-9510 | 1 | auto focus |
| 7 | PAF-DA board, connector | Interface | COP-1317 | 1 | auto focus |
| 8 | W-VIEW GEMINI-2C | Hamamatsu | A12801-10 | 1 | image split |
| 9 | Zoom correction lens unit for W-view-2C | Hamamatsu | A12802-11 | 1 | image split |
| 10 | W-VIEW GEMINI | Hamamatsu | A12801-01 | 2 | image split |
| 11 | ORCA-Fusion BT | Hamamatsu | C15939-20U | 2 | sCMOS camera |
| 12 | Galvanometer Scanner system | Cambridge Tech. | 6220H | 1 | laser angle control |
| 13 | Galvanometer Scanner mount | Riken | Custom-made to connect to a rod | 2 | laser angle control |
| 14 | Function Generator | TEXIO | AFG-3022 | 1 | laser angle control |
| 15 | OBIS LX laser 405 nm×100 mW | Coherent | 87-456 | 1 | laser |
| 16 | OBIS LX laser 488 nm×150 mW | Coherent | 87-461 | 1 | laser |
| 17 | OBIS LS laser 561 nm×100 mW | Coherent | 34-232 | 1 | laser |
| 18 | OBIS LX laser 637 nm×140 mW | Coherent | 87-464 | 1 | laser |
| 19 | OBIS LX/LS 6-Laser Remote | Coherent | 87-475 | 1 | laser power control |
| 20 | Fiber Laser 703 nm×300 mW | MPB Communications | 2RU-VFL-P-300-703-B1R | 1 | laser |
| 21 | Shutter system | OptoSigma | SSH-25RA/SSH-C2B | 3 | shutter for lasers |
| 22 | 5-band dichroic mirror (705) | Chroma | ZT405/488/561/640/705rpc | 1 | dichroic for 4 color imaging |
| 23 | T820dcspxrxt-UF2 | Chroma | T820dcspxrxt-UF2 | 1 | shortpass filter for PAF (830 nm laser) |
| 24 | Filter cube for Nikon Ti2 cube | Chroma | 91032 | 2 | shortpass filter for PAF (830 nm laser) |
| 25 | Bandpass filter (716 nm) for laser | Semorck | FF01-716/40-25 | 1 | Excitation filter for Laser (703 nm) |
| 26 | Bandpass filter (632 nm) for laser | Chroma | ZET642/20x, 25 mm Dia | 1 | Excitation filter for Laser (637 nm) |
| 27 | Bandpass filter (480 nm) for laser | Semorck | FF01-480/17-25 | 1 | Excitation filter for Laser (488 nm) |
| 28 | Dichroic mirror (659 nm) for laser | Semorck | LM01-659-25 | 1 | Dichroic for Lasers 637/703 nm |
| 29 | Dichroic mirror (613 nm) for laser | Semorck | LM01-613-25 | 1 | Dichroic for Lasers 561/637 nm |
| 30 | Dichroic mirror (532 nm) for laser | Chroma | RT532rdc-UF2, 25 mm Dia | 1 | Dichroic for Lasers 488/561 nm |
| 31 | Dichroic mirror (405 nm) for laser | Semorck | Di02-R405-25-D | 1 | Dichroic for Lasers 405/488 nm |
| 32 | Dichroic mirror (640 nm) for image | Semorck | FF640-FDi01-25x36 | 1 | Dichroic in Wview-2C |
| 33 | Dichroic mirror (720 nm) for image | Chroma | T720lpxr-UF1 25.5 x 36 x 1mm | 1 | Dichroic in Wview-1C (long, SF650/SF720) |
| 34 | Bandpass filter (760 nm) for image | Chroma | ET760/50m, 25 mm Dia | 1 | Emission filter in Wview-1C (L-L, SF720) |
| 35 | Bandpass filter (685 nm) for image | Chroma | ET685/50m, 25 mm Dia | 1 | Emission filter in Wview-1C (L-S, SF750) with ET670/50m |
| 36 | Bandpass filter (670 nm) for image | Chroma | ET670/50m, 25 mm Dia | 1 | Emission filter in Wview-1C (L-S, SF750) with ET685/50m |
| 37 | Dichroic mirror (560 nm) for image | Chroma | T560lpxr-UF21, 25.5 x 36 x 2mm | 1 | Dichroic in Wview-1C (short, GFP/TMR) |
| 38 | Bandpass filter (525 nm) for image | Chroma | ET525/50m, 25 mm Dia | 1 | Emission filter in Wview-1C (S-S, GFP) |
| 39 | Bandpass filter (600 nm) for image | Semorck | FF01-600/52-25 | 1 | Emission filter in Wview-1C (S-L, TMR) |
| 40 | 1/2" Achromatic 1/4 Wave plate 350- 800 nm | Thorlabs | AQWP05M | 1 | wave plate |
| 41 | Rotation mount | Thorlabs | RSP1X15/M | 1 | wave plate mount |
| 42 | Post clamp | OptoSigma | PSCA-11.5 | 20 | Fixing of opticsoptics |
| 43 | Post clamp | OptoSigma | RC-10-17 | 20 | Fixing of optics |
| 44 | Post clamp | OptoSigma | RC-10-25 | 20 | Fixing of optics |
| 45 | Post | OptoSigma | PST30 | 20 | Fixing of optics |
| 46 | Post | OptoSigma | PST20 | 20 | Fixing of optics |
| 47 | Rod stand | OptoSigma | RS-12-30 | 20 | Fixing of optics |
| 48 | Rod stand | OptoSigma | RS-20-30 | 20 | Fixing of optics |
| 49 | Stand base | OptoSigma | RCA-M16 | 20 | Fixing of optics |
| 50 | Mirror holder (25 mm) | OptoSigma | MHG-HS25-NL | 6 | Fixing of optics |
| 51 | Mirror holder (25.4 mm) | OptoSigma | MHAN-25.4S | 8 | Fixing of optics |
| 52 | Mirror holder (30 mm) | OptoSigma | MHG-HS30-NL | 8 | Fixing of optics |
| 53 | Mirror 30 mm | OptoSigma | TFAG-30C05-10 | 10 | mirror |
| 54 | Beam expander holder | OptoSigma | KLH-BE-M22H | 5 | Fixing of optics |
| 55 | Beam expander | OptoSigma | LBED3 | 5 | beam expand |
| 56 | Cross cramp | OptoSigma | CCHN-20-20 | 20 | Fixing of optics |
| 57 | Ring for cross cramp | OptoSigma | TR20 | 4 | Fixing of optics |
| 58 | lens holder | OptoSigma | LHF-25S | 1 | filter holder |
| 59 | lens holder | OptoSigma | LHG-25.4 | 4 | filter holder |
| 60 | Rod | OptoSigma | RO-20-500 | 4 | Height Adjustment Platform |
| 61 | Rod | OptoSigma | RO-20-300 | 4 | Height Adjustment Platform |
| 62 | Rod | OptoSigma | RO-20-250-set(4) | 1 | Height Adjustment Platform |
| 63 | Rod | OptoSigma | RO-20-400-set(4) | 1 | Height Adjustment Platform |
| 64 | Rod stand | OptoSigma | RS-6-35 | 1 | Fixing of optics |
| 65 | Breadboad | OptoSigma | OBC-4560-M6 | 1 | Height Adjustment Platform |
| 66 | Height Adjustment Platform | OptoSigma | UP-2030-A430 | 2 | Height Adjustment Platform |
| 67 | iris diaphragm | OptoSigma | IH-12R | 1 | iris |
| 68 | Achromatic lens | OptoSigma | DLB-20-40PM | 1 | Beam expand |
| 69 | Achromatic lens | OptoSigma | DLB-30-150PM | 1 | Beam expand |
| 70 | Achromatic lens | OptoSigma | DLB-30-80PM | 2 | Galvanometer/collimator |
| 71 | Achromatic lens | OptoSigma | DLB-30-100PM | 1 | Galvanometer |
| 72 | Dividing Type Filter Holder Adaptive Element Size Φ15mm | OptoSigma | NDWH-15SRO | 1 | ND filter |
| 73 | ND filter (0.1, 1, 5, 25, 50%) | OptoSigma | AND-15C | 5 | ND filter |
| 74 | Wave plate holder | OptoSigma | PH-30ARS | 1 | Fixing and adjustment of optics |
| 75 | Lens holder with centering adjustment | OptoSigma | LHCM-30 | 3 | Fixing and adjustment of optics |
| 76 | Carriers for Large Optical Rails | OptoSigma | CAA-40LS | 3 | Fixing and adjustment of optics |
| 77 | X axis Aluminum Rack and Pinion Dovetail Stages | OptoSigma | TARA-4025 | 3 | Fixing and adjustment of optics |
| 78 | Adapter Plates | OptoSigma | SP-113N | 3 | Fixing and adjustment of optics |
| 79 | XY-stage | OptoSigma | TSD-802S | 2 | Fixing and adjustment of optics |
| 80 | Magnet base | OptoSigma | MB-L65C-M4 | 2 | Fixing and adjustment of optics |
| 81 | Magnet base | OptoSigma | MB-PH | 2 | Fixing Microscope |
| 82 | Vibration Isolation Systems | OptoSigma | HOA-2010LA | 1 | Base/Vibration Isolation |
| 83 | Silent Air Compressor | OptoSigma | PC-10H-S | 1 | Vibration Isolation |
| 84 | M6 ×22 | Osaka Damashi | 14 mm x 32 | 2 | Fixing of optics |
| 85 | M6 ×22 | Osaka Damashi | 22 mm x 32 | 2 | Fixing of optics |
| 86 | anti-vibration rubber | Hikari | WBG10-10 | 4 | Vibration Isolation |
| 87 | NKB inch Bolt Hexagon Head | Misumi | SNSS-#4-40X3/8 x 10 | 1 | Fixing of optics |
| 88 | Dark room | Morimoto Kasei | MEDR-2200×1200×2000-S1 | 1 | Shading and windproofing |
| 89 | RS232C-USB cable | UGREEN | 20223 | 10 | Connection of PC and control devices |
| 90 | ExComputer BTO02121052401138 | Tsukumo | WA9J-H200/XT | 1 | workstation for system control |
